# DeepBioGS: a hybrid framework for integrating crop growth modelling with genomic prediction through neural networks

**DOI:** 10.64898/2026.05.11.724249

**Authors:** Abdulqader Jighly, Reem Joukhadar, Richard Trethowan, Hans Daetwyler, German Spangenberg

## Abstract

Ensuring global food security under rapid climate change demands accelerated genetic gain and breeding strategies that address complex Genotype-by-Environment (G×E) interactions. Traditional genomic selection models often fail to account for novel or extreme climates.Furthermore, integrating mechanistic crop growth models (CGMs) using traditional Bayesian frameworks to solve this issue presents severe computational bottlenecks. Here, we introduce DeepBioGS, a novel hybrid framework that integrates genomic selection with biophysical growth modelling via a fully differentiable deep learning architecture. DeepBioGS utilises a parameter-prediction multi-layer perceptron to map high-dimensional genomic markers to latent, highly heritable physiological traits (Genotype-Specific Parameters; GSP). These parameters mechanistically predict crop phenology across diverse environments. Using two multi-environment wheat datasets comprising over 6,000 genotypes, DeepBioGS extracted latent traits with near-perfect SNP-based heritability values (0.95-1.00). Crucially, the framework demonstrated superior or comparable predictive accuracy (up to *r*^*2*^ = 0.77) against standard genomic best linear unbiased prediction (GBLUP) and traditional Bayesian CGM-WGP models. Its architecture drastically improved computational scalability by enabling standard backpropagation, effectively bypassing the stochastic sampling limitations of approximate Bayesian methods. Most importantly for climate adaptation, DeepBioGS allowed accurate forecasting of genotype performance in entirely unobserved environmental conditions. By merging the representational power of deep learning with the structural constraints of biophysics, DeepBioGS provides a highly scalable, interpretable tool to navigate G×E interactions, enabling the assessment of cultivars under future climate scenarios, thus optimising crop breeding for a changing global environment.

## Introduction

Ensuring global food security in the face of rapid climate change requires accelerated rates of genetic gain in crop breeding programs. Genomic selection (GS) has revolutionised plant breeding by allowing breeders to predict the performance of their populations based on marker profiles even before they are evaluated in the field (Meuwissen et al. 2001). GS has increased the scale of germplasm evaluation and shortened the duration of breeding cycles, thereby accelerating rates of genetic gain (Heffner et al. 2010). However, the efficiency of traditional GS models is often constrained by their statistical nature. These models typically estimate the effects of genetic markers as random variables within a purely statistical framework, often overlooking the complex biological processes that mediate the relationship between the genome and the phenotype (Technow et al. 2015). These models frequently struggle to account for Genotype-by-Environment (G×E) and Genotype-by-Environment-by-Management (G×E×M) interactions, which are major determinants of phenotypic expression in quantitative traits such as grain yield and phenological development (Messina et al. 2018). Because the effects of markers are estimated based on performance in specific environments, traditional GS models often become less accurate or fail to generalise to novel or extreme environmental conditions where the physiological response of the plant may be non-linear or constrained by non-linear biological processes (Jighly et al. 2026b).

To bridge this gap, researchers have increasingly focused on the integration of Crop Growth Models (CGMs) with GS, creating a unified framework known as CGM-WGP (Technow et al. 2015; Onogi et al. 2016; Messina et al. 2018; Jighly et al. 2023a, 2023b, 2024). CGMs are mechanistic simulations built upon decades of physiological and agronomic research, designed to represent the functional relationships between plant development, resource capture, and environmental variables (Ersoz et al. 2020). Therefore, instead of relying on purely statistical associations as in GS, this approach simulates the physical system by modelling underlying causal mechanisms. This mechanistic physics engine can be used to simulate crop development over time by integrating daily environmental and management inputs to improve the prediction (Chenu et al. 2017). By embedding these biological principles into the predictive process, CGM-WGP attempts to decompose complex traits into their underlying physiological components, which are governed by Genotype-Specific Parameters (GSPs; Washburn et al., 2020). These GSPs are theoretically assumed to be pure genetic variables with heritability equal to 1 (theoretically) and are considered more stable across environments than the complex phenotypes themselves (Heslot et al. 2014). The significance of the CGM-WGP model lies in its ability to unravel G×E interactions by mapping genetic markers directly to these GSPs (Technow et al. 2015). In this framework, the CGM acts as a biological engine that processes environmental inputs and GSPs to simulate the final phenotypes. This approach not only improves the accuracy of predictions in unobserved environments but also enables the prediction of unphenotyped traits and the assessment of cultivars under future climate scenarios (Jighly et al. 2023a).

The practical implementation of traditional Bayesian CGM-WGP frameworks faces several challenges, primarily centred on the computational intensity of parameter estimation and the mathematical complexity of the underlying likelihood functions (Dauda and Lamidi 2026). This forces researchers to rely on Approximate Bayesian Computation (ABC) or Markov Chain Monte Carlo (MCMC) methods, creating a significant computational bottleneck, especially as the number of parameters and their dependencies or correlations increases (Bardenet et al. 2017). In genomic selection contexts, where thousands of markers are mapped to GSPs for which phenotypic data is often unavailable, the probability of randomly sampling a viable parameter set is infinitesimally small (Dauda and Lamidi 2026). This leads to extremely high rejection rates and excessive training times, often requiring massive numbers of iterations to reach convergence (Robert et al. 2011). The slow convergence is further inflated by high linkage disequilibrium, such as that in self-pollinated crops, because highly correlated features in genomic datasets reduce the power of variable selection during the MCMC (Favorov et al. 2005; Kärkkäinen and Sillanpää 2012). Addressing these limitations requires a fundamental rethinking of how mechanistic models are integrated with genomic prediction, moving toward a framework that can leverage the scalability and efficiency of modern deep learning.

Deep learning offers a robust mathematical solution to the bottlenecks inherent in standard CGM-WGP systems. A multi-layered perceptron (MLP) can be implemented to handle the mapping of high-dimensional genomic information to specific physiological traits, which can be used to predict the observed phenotypes through the CGM. Thus, a neural network can learn to estimate latent physiological traits or GSPs by minimising the error between predicted and observed phenotypes using standard backpropagation.

In this study, we developed DeepBioGS, a novel hybrid framework for efficiently integrating genomic selection and biophysical growth modelling using a deep learning architecture to address the limitations of traditional Bayesian methodologies. The system consists of two interconnected, fully differentiable components, a parameter-prediction neural network and a mechanistic physics engine through the CGM. We used a case example here in wheat phenology from a previously published CGM (Christy et al. 2019) to simulate heading (HD) and maturity dates (MD).

## Materials and methods

### Data sets

Two datasets were used to evaluate the accuracy of the DeepBioGS model against other models. The first (Dataset 1) was adopted from Jighly et al. (2023b), which was used to evaluate the CGM-WGP model in the original publication. This data comprised 3,481 genotypes that were phenotyped in Horsham, Victoria, Australia (Lat: −36.71, Long: 142.18) in 11 irrigated and rainfed field trials between years 2012 and 2018. Each field trial comprised between 490 and 1,583 individuals. The second dataset (Dataset 2) comprised 2,570 genotypes phenotyped across 15 normal and heat stressed field trials in three Australian locations: Horsham, Victoria (Lat: −36.71, Long: 142.18; 5 trials); Narrabri, New South Wales (Lat: −30.32, long: 149.83; 8 trials); and Merredin, Western Australia (Lat: −31.48, long: 118.22; 2 trials). The Horsham and Merredin trials had 199 phenotypic records for a core population, while Narrabri trials had between 682 and 1,955 phenotypic records per trial. All field trials had at least two replicates (Joukhadar et al. 2021; Jighly et al. 2026b).

Both datasets were phenotyped for heading date (HD) and maturity date (MD). HD was recorded as the number of days from sowing to 50% emergence of spikes in each plot. MD was recorded on the day when 90% of the plot reached physiological maturity defined as stage 90 of the standard Zadoks score (Zadoks et al. 1974). Dataset 2 in Merredin only had HD data without MD in both trials. The two datasets were genotyped with 90K Infinium SNP chip (Wang et al. 2014) and only SNPs with minimum call rate >50% and minor allele frequency >5% (34,552 SNPs for the first dataset and 41,666 SNPs for the second dataset) were used for all genomic analyses.

### Overview of the DeepBioGS framework

We implemented a hybrid Crop Growth Model - genomic selection framework using a deep learning architecture (Figure 1). The system consists of two interconnected, fully differentiable components: a parameter-prediction neural network (Multilayer Perceptron) and a mechanistic physics engine (the Crop Growth Model). The neural network translates genotypic marker profiles into latent cultivar-specific physiological parameters, which are then passed into the crop growth model. The crop growth model uses daily weather variables to predict the observed heading date (HD) and maturity date (MD). The neural network learned to estimate the latent physiological traits by minimising the error between the predicted and observed phenological dates obtained by applying the CGM using standard backpropagation (Figure 1). The model was run for 50 random replicates and in each one, half of the population was randomly selected for validation, and the other half was used to train the model.

**Figure 1.**
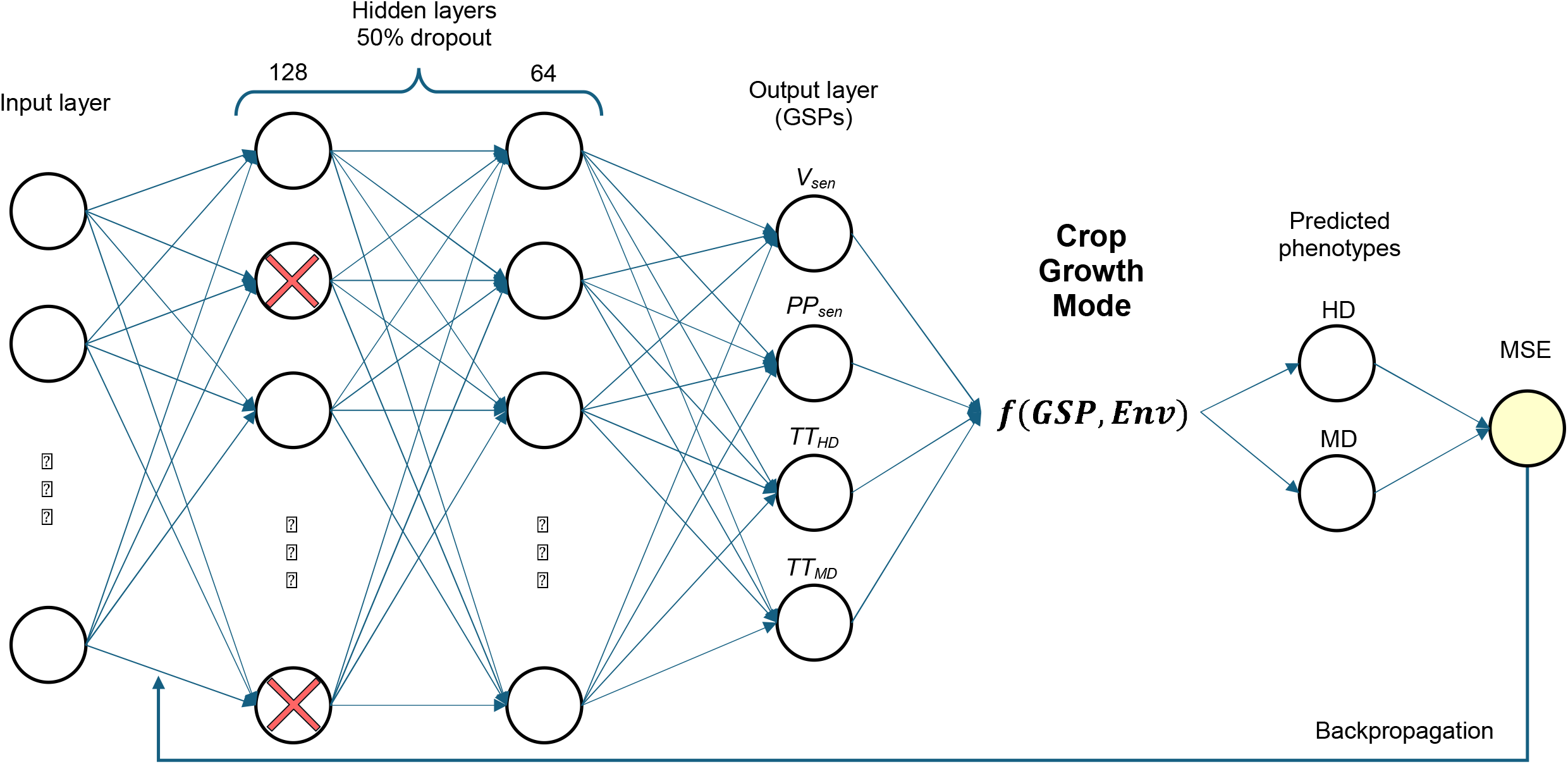
The architecture of the DeepBioGS model

#### Genotype-to-GSP multi-layer perceptron

To map high-dimensional genomic information to physiological traits, a multi-layer perceptron (MLP) was implemented (Popescu et al. 2009), using the Python TensorFlow package (Abadi et al. 2015). The input layer consisted of scaled genomic markers. The architecture comprised:

- A network to receive the genotype marker profiles as input. This is followed by two hidden dense layers containing 128 and 64 neurons, respectively, both utilising Rectified Linear Unit (ReLU) activation functions. These functions allow the network to learn complex, non-linear patterns in the genetic data.
- A dropout layer with a rate of 50%. This was applied between the two hidden layers during training to prevent overfitting of the genomic inputs.
- A final output layer consisting of four nodes with a sigmoid activation function.

The final output layer of the MLP consists of four nodes with a sigmoid activation function, which predicted the latent Genotype-Specific Parameters (GSPs) for each genotype. These parameters are essential biological traits that dictated how the crop responds to environmental cues including:

- *v*_*sen*_: Vernalisation sensitivity, which determines the crop’s requirement for cold temperatures to initiate the transition from vegetative to reproductive growth.
- *pp*_*sen*_: Photoperiod sensitivity, representing the plant’s developmental response to day length.
- *TT*_*HD*_: Target thermal time to heading, the accumulated warmth required to reach the heading stage.
- *TT*_*MD*_: Additional target thermal time required from heading to maturity.

#### Mechanistic phenology crop growth model

The predicted GSPs are passed directly into the second component: a mechanistic physics engine that simulates the daily phenological development of the crop based on environmental variables. This engine was built upon the phenology model described by Christy et al. (2019), which integrated daily temperature and photoperiod data to track development progress.

For each simulated day, the engine calculated the effective daily thermal time accumulation (*TTdaily*) as:

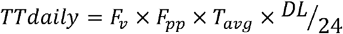

Where *T*_*avg*_ is the daily average temperature, and *DL* is the day length in hours. The modifiers *F*_*v*_(vernalisation factor) and *F*_*pp*_ (photoperiod factor) restrict development based on the environment and the genotype’s specific sensitivities:

##### Vernalisation Modifier (*F*_*v*_)

*F*_*v*_ is derived from the cumulative cold experience of the crop with value starts at 0 and reach 1 at floral initiation, beyond which, its value will always equal to one. It can be calculated for each day as:

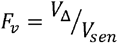

Where *v*_*Δ*_ is the cumulative vernalisation time experienced by the crop as:

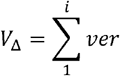

Where *ver* represents the accumulation of vernalisation requirements, which equal 1 in cold days with *T*_*avg*_ between 2 and 9 °C. The value of *ver* decreases linearly outside this optimal cold range to reach a value of 0 when the *T*_*avg*_ becomes lower than −4 °C or higher than 15 °C (White et al., 2008; Christy et al., 2019).

##### Photoperiod Modifier (*F*_*pp*_)

*F*_*pp*_ensures that the rate of development is biologically responsive to the environmental context and its values range between 0 and 1 and for each day and is calculated as:

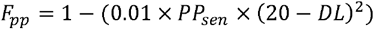

The cumulative thermal time *TT* up to any given day (*n*) is the sum of the daily effective thermal times as:

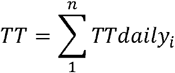

Simulated HD was assumed on the day when *TT* becomes equal to or larger than *TT*_*HD*_; while simulated MD was assumed on the day when *TT* becomes equal to or larger than *TT*_*HD*_ + *TT*_*MD*_

### Model training

The framework was trained to jointly predict HD and MD with the following assumptions:

- Loss Function: The network minimised a weighted Mean Squared Error (MSE) between the observed and predicted days to heading and maturity. The loss was computed exclusively over valid (non-missing) observations. To account for data imbalance across environments, the loss was weighted based on the frequency of each genotype’s occurrence in the dataset.
- Optimisation: The network weights were optimised using the Adam optimiser with a learning rate of 0.001.
- Training duration: The model was trained for 200 epochs. This number was selected after testing multiple values for which the aim was to select a total number of epochs that is long enough to ensure proper divergence but not too long to avoid overfitting. Generally, overfitting started to happen after around 500 iterations.

To quantify the predictive uncertainty in predicting the GSPs, we employed a probabilistic deep learning framework (Patel et al. 2015; Gal and Ghahramani 2016). Rather than relying on the point estimates, this approach captured the model’s confidence by treating the prediction as a distribution. Uncertainty estimation was performed using the finalised model after the training phase; while the learned weights remained fixed, dropout was kept active during the inference stage to introduce stochasticity. We implemented 500 Monte Carlo (MC) sampling iterations after completing the model training to approximate the predictive posterior distribution. This stochastic sampling allowed calculation of the mean GSP alongside its associated variance, providing a robust measure of uncertainty.

### Benchmark

DeepBioGS was compared to the Genomic Best Linear Unbiased Prediction (GBLUP) model without (EG) and with consideration of genotype by environment interactions (G×E). The general equation for both implementations was:

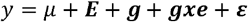

Where, y is the phenotypic record (HD or MD), μ is the intercept, E is the environment effect which follows normal distribution 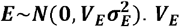 was calculated by multiplying *z*_*E*_ (n × e) incidence matrix with its transpose, allowing allocation of n genotypes to e environments; **g** represents the genotypic effects following a normal distribution 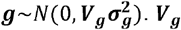 was calculated as ***V***_***g***_***=Z***_***g***_***GZ***^***’***^_***g***_, in which *z*_***g***_ (m × n) was the incidence matrix allocating phenotypic records to genotypes, with m representing the number of variants. **G** (n × n) is the genomic relatedness matrix (method 1 in: VanRaden 2008). ε is the independent and identically distributed residuals, 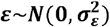.

The ***gxe*** represents the genotype by environment interaction term in the equation. For the model that does not consider the interaction (EG), ***gxe*=**0. On the other hand, for the G×E model, 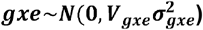, where ***v***_*gxe*_ = ***v***_*g*_⨀***v***_*E*_, and ⨀ represents the cell-to-cell product (Hadamard product). The model was fitted using the R package BGLR (Pérez and de Los Campos, 2014). Both models were conducted with a total of 20,000 iterations for which the first 10,000 iterations were discarded as burn in.

### Cross-validation

The predictive performance of the models was evaluated using two distinct cross-validation strategies. The first strategy, CV1, assessed the model’s ability to predict the performance of individuals that had never been phenotyped. For this approach, the population was randomly partitioned into training and validation sets of equal size (50% each). The second strategy involved masking a subset of the environments to be predicted to evaluate the model’s capability to generalise to entirely unobserved environmental conditions. For this strategy, two schemes were employed. First, a leave-one-environment-out approach (OneOut) to evaluate the model performance in predicting new environments using a large representative reference. Under this scheme, each environment was iteratively excluded from the training set and used as the validation population. Secondly, a single-environment reference approach (Single) was used to evaluate prediction of new environments when the reference was limited. Under this scheme, each environment was iteratively used alone for training, and the remaining environments were used for validation. It is important to note that the GBLUP-G×E model was inapplicable to the second strategy, as it generates environment-specific genomic estimated breeding values (GEBVs) that cannot be directly generalised to new sites and such indirect generalisation is less accurate.

For each evaluated scenario, 50 random replicates were performed for each model and trait combination. Prediction accuracy was quantified as the Pearson correlation coefficient between the observed phenotypes and the predicted GEBVs of the validation individuals. The mean accuracy across all replicates was reported for more robust estimation. For Dataset 1, the results were benchmarked against the prediction accuracies reported by Jighly et al. (2023b) for the traditional Bayesian CGM-WGP model. This direct comparison was facilitated by using the same dataset across both studies; this benchmarking approach was necessary as the original Bayesian implementation was not locally accessible for direct re-running during this study. Student’s t-test was used across the replicates to declare significant differences in prediction accuracy among evaluated scenarios for both HD and MD.

### Narrow sense heritability, and genetic correlation

The sampled GSPs within both datasets were analysed with MTG2 software (Lee and Van der Werf 2016) to calculate the SNP-based heritability (which is equivalent to the narrow-sense heritability) with the software’s default parameters. The same software was supposed to estimate the genetic correlation among the GSPs sampled from each dataset. However, because the SNP-based heritability values for all sampled GSPs were around one, this analysis became technically unreliable. Having a near zero error variance value in both traits under investigation could lead to singular error matrix that would disturb the convergence of the model.

## Results

The analysis of the sampled GSPs with the DeepBioGS model revealed exceptionally high SNP-based heritability (*h*^*2*^) across both evaluated datasets, validating the theoretical assumption of highly stable genetic variables (Table 1). Specifically, in the first dataset, the heritability estimates reached 1.00 for both *v*_*sen*_ and *pp*_*sen*_, while the target *TT*_*HD*_ and additional target *TT*_*MD*_ exhibited heritability values of 0.97 and 0.95, respectively. These *h*^*2*^ values are much higher than previously reported by Jighly et al. (2023b), who calculated *h*^*2*^ for the same GSPs estimated with the Bayesian CGM-WGP model on the same dataset, which had an average *h*^*2*^ value of 0.76. Similarly, the second dataset demonstrated near-perfect heritability scores, with *v*_*sen*_ calculated at 0.98, *pp*_*sen*_ and *TT*_*HD*_ both reaching 1.00, and *TT*_*MD*_ at 0.99.These near-maximum values approaching 1.00 indicate a minimal error variance for the latent physiological traits extracted by the model’s neural network component across varying environmental conditions.

**Table 1.** SNP-based heritability (*h*^*2*^) of the four genotype-specific parameters (GSPs) in both datasets.

| **GSP** | **$h^2$ dataset1** | **$h^2$ dataset2** |
| --- | --- | --- |
| $V_{sen}$ | 1.00 | 0.98 |
| $PP_{sen}$ | 1.00 | 1.00 |
| $TT_{HD}$ | 0.97 | 1.00 |
| $TT_{MD}$ | 0.95 | 0.99 |

In the evaluation of predictive performance using the dataset1, DeepBioGS consistently demonstrated robust accuracy across multiple CV scenarios that was comparable to the accuracies of the standard baseline models (Table 2). Under the single-environment reference scheme (Single), DeepBioGS effectively generalised from limited reference data, achieving a prediction accuracy of 0.76 and an RMSE of 12.2 days for HD, alongside a prediction accuracy of 0.66 and RMSE of 11.8 days for MD. This significantly outperformed the baseline EG model, which yielded lower accuracies of 0.70 for HD and 0.59 for MD in the same scenario. When predicting entirely unobserved environments via the OneOut approach, DeepBioGS reached a prediction accuracy of 0.77 for HD and 0.69 for MD, matching the reported HD accuracy of the traditional Bayesian CGM-WGP model and slightly, but not significantly, exceeding its MD accuracy (*r*^*2*^ = 0.67). Furthermore, when predicting the performance of non-reference individuals (CV1), DeepBioGS maintained high predictive accuracies of 0.77 for HD and 0.69 for MD with low respective RMSE values of 7.2 and 4.5 days, surpassing both the standard EG model (*r*^*2*^ = 0.75 for HD; 0.67 for MD) and the G×E model (*r*^*2*^ = 0.74 for HD; 0.67 for MD), but the differences were not statistically significant.

**Table 2.** Root mean squared error (RMSE) and prediction accuracy (*r*^*2*^) for dataset1 using different models and cross-validation (CV) scenarios. Single: Single reference environment and remaining environments for validation; OneOut: Single validation environment and remaining environments were used in the reference; CV1: all environments in the reference, predicting non-reference individuals

| Model | CV | HD RMSE | HD $r^2$ | MD RMSE | MD $r^2$ |
| --- | --- | --- | --- | --- | --- |
| **EG** | **Single** | NA <sup>a</sup> | 0.70 | NA <sup>a</sup> | 0.59 |
|  | **OneOut** | NA <sup>a</sup> | 0.75 | NA <sup>a</sup> | 0.67 |
|  | **CV1** | 7.8 | 0.75 | 4.6 | 0.67 |
| **G×E** | **CV1** | 6.9 | 0.74 | 4.1 | 0.67 |
| **CGM-WGP** <sup>b</sup> | **Single** | NA <sup>c</sup> | 0.70 | NA <sup>c</sup> | 0.59 |
|  | **OneOut** | NA <sup>c</sup> | 0.77 | NA <sup>c</sup> | 0.67 |
|  | **CV1** | NA <sup>c</sup> | 0.77 | NA <sup>c</sup> | 0.67 |
| **DeepBioGS** | **Single** | 12.2 | 0.76 | 11.8 | 0.66 |
|  | **OneOut** | 7.7 | 0.77 | 4.9 | 0.69 |
|  | **CV1** | 7.2 | 0.77 | 4.5 | 0.69 |

<sup>a</sup> RMSE calculation is not applicable to EG model for single and OneOut trait analysis as the predicted values are centred around the average of the reference single trial leading to RMSE>60 days when predicting other environments in extreme cases
<sup>b</sup> Results as reported in Jighly et al. (2023b) who worked on the same dataset.
<sup>c</sup> RMSE for CGM-WGP model reported in Jighly et al. (2023b) were calculated using a subset of the lines, which was different CV strategy compared to the methods described here

The robust predictive capability of DeepBioGS was further confirmed in dataset2, where it maintained similar or slightly higher accuracies despite slightly lower baseline accuracies across most of the evaluated models for HD and MD (Table 3). In the Single CV scenario, DeepBioGS performed comparably to the standalone EG model, achieving an *r*^*2*^ of 0.54 (RMSE = 9.1 days) for HD and 0.53 (RMSE = 6.9 days) for MD. However, the framework demonstrated stronger generalisation capabilities to novel environments in the OneOut scenario, reaching an *r*^*2*^ of 0.57 for HD and 0.60 for MD, thereby outperforming the standard EG model which only achieved accuracies of 0.53 and 0.58 for the respective traits. The improvement was only statistically significant for HD. Finally, under the CV1 scheme predicting unphenotyped individuals, DeepBioGS yielded the highest overall predictive accuracies for this dataset at 0.57 for HD and 0.60 for MD. In comparison, the G×E model achieved slightly lower HD accuracy (0.56) while matching the MD accuracy (0.60), and the standard EG model lagged with accuracies of 0.54 for HD and 0.58 for MD. However, all these differences in prediction accuracy were not significant.

**Table 3.** Root mean squared error (RMSE) and prediction accuracy (*r*^*2*^) for dataset2 using different models and cross-validation (CV) scenarios. Single: Single reference environment and remaining environments for validation; OneOut: Single validation environment and remaining environments were used in the reference; CV1: all environments in the reference, predicting non-reference individuals

| Model | CV | HD RMSE | HD $r^2$ | MD RMSE | MD $r^2$ |
| --- | --- | --- | --- | --- | --- |
| **EG** | **Single** | NA <sup>a</sup> | 0.53 | NA <sup>a</sup> | 0.54 |
|  | **OneOut** | NA <sup>a</sup> | 0.53 | NA <sup>a</sup> | 0.58 |
|  | **CV1** | 3.2 | 0.54 | 2.7 | 0.58 |
| **GxE** | **CV1** | 2.8 | 0.56 | 2.2 | 0.60 |
| **DeepBioGS** | **Single** | 9.1 | 0.54 | 6.9 | 0.53 |
|  | **OneOut** | 4.1 | 0.57 | 3.3 | 0.60 |
|  | **CV1** | 3.0 | 0.57 | 2.5 | 0.60 |

<sup>a</sup> RMSE calculation is not applicable to EG model for single and OneOut trait analysis as the predicted values are centred around the average of the reference single trial leading to RMSE>60 days when predicting other environments in extreme cases

DeepBioGS showed improved characteristics over the three other models tested (EG, G×E and Bayesian CGM-WGP). The model exhibited significantly enhanced predictive capacity compared to conventional Bayesian CGM-WGP approaches, particularly when reference data was restricted to a single environment. While the G×E model produced slightly lower RMSE for non-reference individuals (CV1) across both datasets, its practical application was constrained by inherent inability to generalise to environments unobserved in the reference set. Similarly, although the EG model allowed for generalisation and can achieve comparable accuracy in certain contexts, it failed to provide accurate phenotypic estimations in novel environments. Because EG model predictions are centred on the reference trial average, the model cannot approximate the arithmetic mean of unobserved environments, often resulting in prohibitive RMSE values exceeding 60 days. In contrast, DeepBioGS avoids these limitations by utilising a differentiable architecture to directly model the mechanistic interactions between genetic parameters and environmental factors, allowing for more robust predictions of final heading and maturity dates.

## Discussion

The development of DeepBioGS represents a strategic move toward a new generation of intelligent breeding tools that reduce the gap between biological knowledge and computational efficiency. By integrating a parameter prediction neural network with a fully differentiable mechanistic physics engine, this hybrid framework addresses the long-standing bottlenecks of traditional genomic selection and environmental modelling. The primary significance of this approach is its ability to transform complex biological simulations into a single, differentiable computational graph that can be optimised using modern deep learning algorithms. The results showed that this structure needed less extensive computational and training data in addition to mitigating the equifinality problem.

The evaluation of DeepBioGS across varied environmental conditions highlighted its capacity to dissect complex G×E×M interactions. The model successfully extracted latent traits with near-perfect heritability values (>0.95) over the four GSPs. GSPs should theoretically be 100% heritable. Sampling GSPs with such high heritability does not mean that DeepBioGS captured the whole genetic variance of the latent traits, instead it shows that the model captured the whole genetic variance that can be explained by the input SNPs given the CGM. Therefore, introducing more input data, e.g. omics or input phenomics data, could allow sampling of wider genetic variance for the GSPs, while the heritability remains high. This interpretability provides breeders with actionable insights into the specific traits driving a cultivar’s performance. Consequently, breeding programs can precisely identify which physiological sensitivities confer robustness under heat stress and target those specific latent traits to stabilise crop development in shifting macroclimates.

The following sections provide a detailed comparative analysis of DeepBioGS against three primary methodologies, traditional standalone CGMs, standalone deep learning models, and conventional Bayesian CGM-WGP solutions (Table 4). These comparisons highlight how the framework leverages the scalability of AI while remaining anchored in the universal biophysical processes that govern crop development.

**Table 4.** Detailed comparisons among traditional CGM, GS through linear and deep learning models, Bayesian CGM-WGP, and DeepBioGS.

| **Feature** | **Traditional CGM** | **Standalone GS and Deep Learning** | **Bayesian CGM-WGP** | **DeepBioGS** |
| --- | --- | --- | --- | --- |
| **Optimisation Method** | Stochastic search | Standard backpropagation | Stochastic search given prior distributions | Backpropagation through the biophysical model |
| **Scalability** | Cultivar-specific | Highly scalable | Exponentially slower with growing data | Highly scalable |
| **Data Requirements** | Very high, on the cultivar level | Medium | Medium | Low |
| **Interpretability** | High; GSPs have direct physiological meaning | Low | High; GSPs have direct physiological meaning | High; GSPs have direct physiological meaning |
| **Gene-interaction** | Not applicable | Captures both linear and non-linear epistasis | Primarily linear assumptions | Captures both linear and non-linear epistasis |
| **Generalisation** | Good only for cultivars with large multi-environmental data | Poor on novel/extreme environments | Robust; anchored in biophysical processes | Robust; anchored in biophysical processes |
| **Technical difficulties <sup>a</sup>** | High equifinality | Overfitting (mainly for DL) | Extensive parameter tuning; Equifinality | Equifinality (less than CGM-WGP) |
| **Computational requirements** | Medium | Low (DL) Medium (Linear GS models) | Very high | Low |
| **Accuracy** | Low | Medium-high | High | High |

<sup>a</sup> Assuming the base CGM is already developed

### Advantages of DeepBioGS over traditional CGM

While traditional CGMs excel in interpretability, where GSPs provide clear physiological insights, they are fundamentally limited by a population-blind architecture (Jighly et al. 2026b). Because these models do not integrate genomic markers or relatedness information, they cannot exploit shared genetic information across a population, requiring high-density phenotypic data for every specific cultivar to achieve accuracy (Ogoni et al. 2016). This cultivar-specific focus makes them unscalable for large-scale breeding programs where thousands of new lines must be evaluated simultaneously, therefore, their applications are limited to established cultivars with data collected across diverse environments (Andrade et al. 2005). It was previously discussed that the main advantage of the Bayesian CGM-WGP over conventional CGM is the transformation of the prediction from individual to population level, which mitigates the equifinality problem due to the correlation between GSPs (Jighly et al. 2023a). Furthermore, the optimisation process in traditional CGMs is characterised by a stochastic slow search that relies on trial-and-error sampling rather than efficient gradient-based methods, making the calibration process computationally demanding (Peng et al. 2020). Crucially, standalone CGMs lack the mechanism to model gene-to-trait interactions, as they do not map genomic profiles to physiological traits, making them unable to account for additive and complex epistatic genetic architectures (Technow et al. 2015). Consequently, these models generalise well only when provided with large multi-environmental datasets for specific, well-characterised varieties, leaving a significant gap in the ability to predict the performance of new, unphenotyped genotypes across diverse environments.

### Advantages of DeepBioGS over standalone deep learning

In DeepBioGS, the neural network acts as a flexible parameter estimator, while the crop growth model acts as a physics engine that enforces biological and physical constraints (Faure et al. 2023). This hybrid architecture offers several unique advantages over standalone deep learning. Firstly, it reduces data requirements. Because the crop growth model incorporates established biological laws (e.g., thermal time accumulation, photoperiodic response), the neural network does not have to learn these fundamental relationships from the data. This allows accurate predictions even with the limited training population sizes typically found in plant breeding (Faure et al. 2023). Secondly, it enforces structural regularisation. The mechanistic equations act as a natural bottleneck that regularises the neural network, preventing it from producing biologically implausible outputs and reducing the risk of overfitting (Stock et al. 2024). Thirdly, it improves generalisation. By anchoring the prediction in universal biophysical processes, hybrid models are more robust when extrapolating to novel or extreme environmental conditions that are not represented in the training set. Lastly, DeepBioGS has much improved interpretability. While pure deep learning models are often black boxes, hybrid models maintain high interpretability because their internal parameters (GSPs) have direct physiological significance (Miranda et al. 2024). This provides breeders with actionable insights into the specific traits driving a cultivar’s performance.

### Advantages of DeepBioGS over conventional Bayesian CGM-WGP solution

DeepBioGS represents a significant computational and methodological evolution over traditional Bayesian CGM-WGP models by replacing the less optimal stochastic searches and trial-and-error sampling with a fully differentiable architecture that utilises backpropagation through the model graph. This transition allows a drastic increase in training speed and faster convergence (Sunnåker et al. 2013). While traditional Bayesian approaches face exponential slowdowns when dealing with high-dimensional data, DeepBioGS is linearly scalable and leverages GPU acceleration, making it feasible to utilise high-density, full-genome marker sets. Furthermore, by moving away from likelihood-free approximations toward the direct minimisation of phenotypic error, the framework achieves more stable parameter estimation and improved predictive accuracy (Faure et al. 2023). These technical improvements are supported by the hard coding of physical laws into the engine, which enforces biological plausibility more effectively than the weak and flexible distributions used in traditional Bayesian models. Ultimately, while traditional methods are often limited by linear assumptions, DeepBioGS can capture both linear and non-linear epistatic interactions, providing a much deeper understanding of complex genetic architectures.

The most significant technological advancement for DeepBioGS is the emergence of differentiable simulators. Modern machine learning frameworks utilise automatic differentiation to compute gradients of complex functions defined in the model (Hamazaki et al. 2025). By reimplementing the mechanistic equations of a crop growth model within these frameworks, the entire system, from genomic input to phenotypic output, becomes a single, differentiable computational graph. This differentiability solves the model by enabling the use of backpropagation and gradient-based optimisers like Adam. Instead of stochastically sampling the parameter space and rejecting unsuccessful simulations, the model can efficiently compute exactly how to adjust the marker effects and physiological parameters to minimise the prediction error (Hamazaki et al. 2025). This leads to orders-of-magnitude faster training times and allows the model to scale to datasets with hundreds of thousands of markers and thousands of environments, making CGM-WGP practical for real-world breeding applications.

Deep learning models have demonstrated superior performance in identifying gene-gene interactions such as epistasis (Montesinos-López et al. 2021). By using a neural network as a parameter-prediction component, the DeepBioGS framework can leverage this representational power to estimate GSPs more accurately than traditional Bayesian regression models and can capture complex interaction patterns among genetic markers (Perelygin et al. 2025). This data-driven approach allows the model to learn the optimal mapping from markers to physiology directly from the data, without the need for manual feature engineering or restrictive assumptions about genetic architecture.

## Conclusion

By combining the power of deep learning with the structural constraints of mechanistic physiological models, DeepBioGS provides a robust solution to the confounding effects of G×E×M in genomic selection. This model allows the precise estimation of latent physiological traits directly from genotypic data, offering a level of interpretability and scalability that traditional GS models cannot match. The development of DeepBioGS thus represents a shift toward a new generation of intelligent breeding tools that are both biologically grounded and computationally optimised for the demands of modern agriculture. Looking forward, the DeepBioGS framework offers a highly flexible architecture with significant potential for expansion. While this study focused on phenology, the system can be extended to incorporate full crop growth models to predict complex traits like grain yield, biomass accumulation and protein content across diverse environmental scenarios. Furthermore, its modular design allows rapid adaptation to a wide range of crop species beyond wheat, enabling the integration of various mechanistic equations tailored to specific biological structures. Future developments of the model could also incorporate high-throughput phenotyping and daily collected data and remote sensing inputs to further refine the estimation of latent physiological parameters in real-time. On the predictor’s side, multi-omics data beyond genotypic data can be also integrated to achieve higher prediction accuracy.

Ensuring global food security in the face of rapid climate change inherently requires accelerating the rate of genetic gain within crop breeding programs. Historically, the integration of crop growth models with genomic selection via traditional Bayesian frameworks has faced severe challenges due to the computational intensity of parameter estimation and the reliance on MCMC methods. DeepBioGS overcomes this critical computational bottleneck by utilising a fully differentiable architecture that employs backpropagation through the model graph. This structural shift drastically increases training speeds, enables linear scalability through GPU acceleration, and comfortably handles high-density, full-genome marker sets. Furthermore, this architecture effectively mitigates the equifinality problem often associated with complex biological modelling. By providing a computationally optimised and biologically grounded tool, DeepBioGS grants breeders the speed necessary to rapidly evaluate vast populations under projected climatic stresses. Ultimately, bridging the gap between genomic potentials and environmental factors paves the way for the rapid development of more resilient and productive agricultural systems equipped for future scenarios with increasing climatic variability

## Conflict of Interest

The authors declare that they have no conflict of interest.

## Acknowledgements

The authors would like to thank Qingdao Agricultural University.

## Data availability

The data used in the present study was previously published in Jighly et al. (2023b); and Jighly et al. (2026a)

## Author contributions

AJ: developed the model; AJ, RJ, GS: planned the study; AJ, RJ: analysed the data; RT: provided the data; AJ, RJ: wrote the first draft; HD, RT, GS: revised the first draft and provided critical comments to improve the content of the manuscript.

## Funding

The study was funded through a grant from Qingdao Agricultural University (GS), China

## Notes

### Competing Interest Statement

The authors have declared no competing interest.

